# Essential Regression - a generalizable framework for inferring causal latent factors from multi-omic human datasets

**DOI:** 10.1101/2021.05.03.442513

**Authors:** Xin Bing, Tyler Lovelace, Florentina Bunea, Marten Wegkamp, Harinder Singh, Panayiotis V Benos, Jishnu Das

**Affiliations:** Department of Statistics and Data Science, Cornell University, Ithaca, NY, USA; Department of Computational & Systems Biology, University of Pittsburgh, Pittsburgh, PA, USA; Joint CMU-Pitt PhD Program in Computational Biology, Pittsburgh, PA, USA; Department of Mathematics, Cornell University, Ithaca, NY, USA; Center for Systems Immunology, Departments of Immunology and Computational & Systems Biology, University of Pittsburgh, Pittsburgh, PA, USA

## Abstract

High-dimensional cellular and molecular profiling of human samples highlights the need for analytical approaches that can integrate multi-omic datasets to generate predictive biomarkers and prioritized causal inferences. Current methods are limited by high dimensionality of the combined datasets, the differences in their data distributions and their integration to infer causal relationships. Here we present Essential Regression (ER), an interpretable machine learning approach for high-dimensional multi-omic datasets, that addresses these problems by identifying latent factors and their likely cause-effect relationships with the system-wide outcome/properties of interest. ER is a novel data-distribution-free latent-factor regression model that integrates multi-omic datasets and identifies latent factors significantly associated with an outcome. ER outperforms a range of state-of-the-art methods in terms of prediction performance on simulated datasets. ER can be coupled with probabilistic graphical modeling thereby strengthening the causal inferences. ER generates novel cellular and molecular predictions, using multi-omic human systems immunology datasets, pertaining to immunosenescence and immune dysregulation.

## Introduction

Over the last decade, genomic, proteomic, metabolomic and other technologies for generating deep molecular profiles of tissues and cells from model organisms or humans have rapidly expanded (1–4). However, the explosion in data, especially from a range of such ‘omic technologies has not been coupled to a proportional increase in our understanding of the underlying causal mechanisms. Existing analytical approaches have primarily focused on individual “omic” datasets with relatively few attempts at integration of multi-omic datasets. In either case, we (5–9) and others (10–12) have primarily emphasized the delineation of predictive biomarkers (Fig. 1a). A key focus of these efforts has been to overcome the “curse of dimensionality” (very large number of variables being measured in relation to a comparatively low number of samples) and the multiplicity of predictive signatures due to multi-collinear data i.e., large correlated sets of variables. While there are several methods for reliably uncovering predictive markers from high dimensional data, none of these analyze cause-effect relationships in relation the outcomes/outputs of interest. This in turn has hampered efforts to undertake perturbative/translational experiments and/or clinical investigations that can test a functionally prioritized set of hypotheses generated by the large datasets.

**Figure 1.**
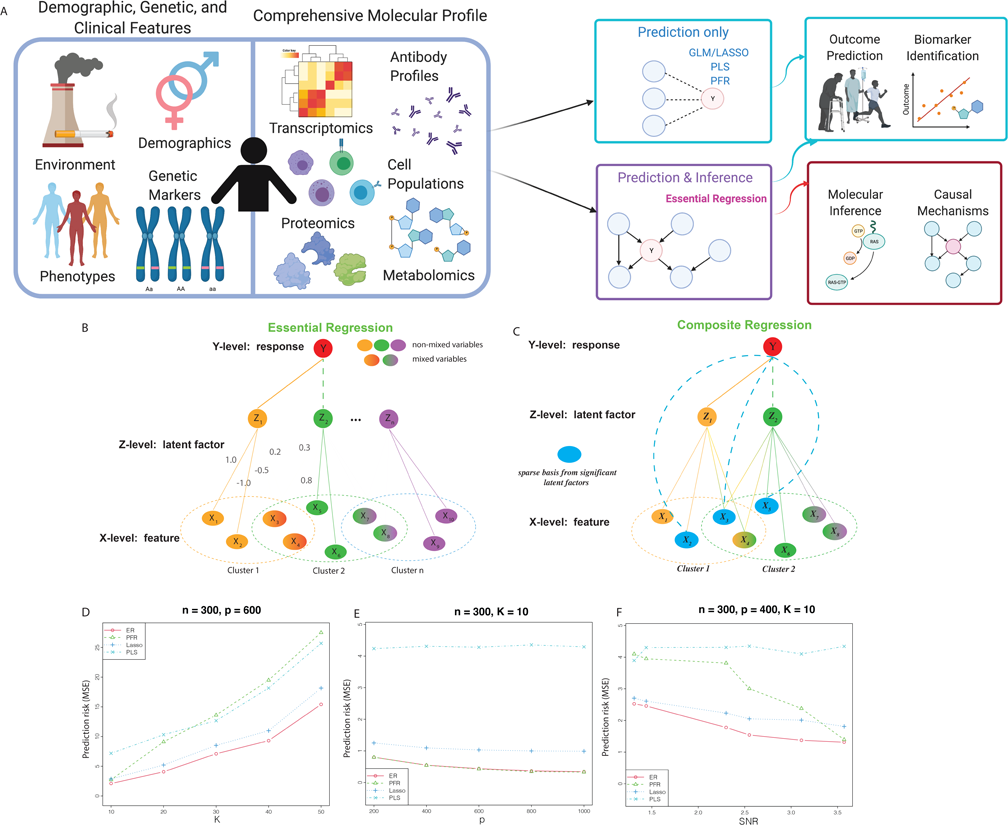
Essential Regression – a novel interpretable amchine learning approach to uncover causal latent factors from high-dimensional multi-omic datasets. a) Schematic illustrating the different kinds of multi-omic datasets typically used in systems analyses and the key advantages of the methods introduced in this study over existing approaches. b) Schematic summarizing the steps in ER. c) Schematic summarizing the steps in Composite Regression. d-f) Comparison of the predictive performance of PLS, PFR, LASSO and ER on simulated datasets across a range of parameter settings.

In addition to the high dimensionality of datasets at any given scale of organization (e.g., cellular, molecular), biological systems, particularly humans, manifest extreme complexity in terms of numbers of molecular components, their interaction rules as well as their hierarchical scales of organization that include macromolecular complexes/condensates, organelles, cells, tissues and organs. Each scale of organization in such a complex system has components and interaction rules that are unique to its level of organization. Thus, predicting changes in properties or behaviors of the system based on measuring components that are operating at different scales of organization represents a formidable challenge. Methods that make assumptions regarding data-generating mechanisms typically perform poorly at multi-scale integration as there are key differences in data distributions at each scale of organization.

We propose a novel framework, Essential Regression (ER), to address these key challenges and limitations of existing approaches by focusing on latent factors rather than observables in high dimensional datasets that are significantly associated with a system wide-property or outcome that is of interest (Fig. 1a). Critically, ER makes no assumptions regarding the underlying data distributions, enabling principled integration of multi-omic datasets. Further, ER uses regression on the latent factors rather than the observables, a novel statistical framework that comes with rigorous guarantees. Overall, our analytical framework derives causal latent factors from thousands of variables from multi-omics datasets across various scales of biological organization (Fig. 1a). After identifying significant latent factors, ER can be coupled with causal graphical model analyses to examine the connectivity of these factors to the system-wide property or outcome of interest. In so doing ER generates a high confidence and prioritized set of latent factors comprised of known observables that are most proximal in the causal graph network to the system property/outcome of interest. We note that while causal discovery approaches have become popular over the last two decades, they have been confined to low-dimensional datasets due to the associated computational complexity. ER overcomes this fundamental conceptual limitation by first identifying latent factors from the observables (which achieves an inherent dimensionality reduction), and then identifying which latent factors are causally linked to the outcome/system wide property of interest.

By analyzing both simulated and real-world human immunological multi-omic datasets, we demonstrate that ER and the subsequent causal graphical model analyses significantly outperform a wide range of state-of-the-art approaches in predicting outcomes and provides multi-scale inferences not afforded by the existing methods. The novel predictions are corroborated by biological findings in model systems.

## Results

### Essential Regression – a novel data-distribution-free statistical regression framework for inferring causal latent factors

We present Essential Regression (<u>ER</u>), a novel data-distribution-free latent factor regression approach that integrates high-dimensional multi-omic datasets and identifies latent factors that are significantly associated with a system property/outcome (Fig. 1b, Methods, Supplementary Note 1). ER is a paradigm-altering concept in regression analysis for high dimensional datasets i.e., datasets where the number of features exceeds the number of samples. Existing regression methods use techniques including regularization (e.g., L1 regularization – LASSO(13), L1 + L2 regularization – Elastic Net), bootstrap aggregation (e.g., random forest(14)) or the incorporation pre-specified group structure (e.g., group LASSO) to avoid overfitting. However, biomarkers/features identified using these approaches are merely predictive/correlative and may have no connection with the underlying mechanisms driving the system property/outcome of interest. ER, on the other hand uses a two-step approach that allows for the identification of latent factors that can be used to infer causal structures underlying the system property/outcome. ER first finds latent factors in a data-dependent fashion, without the need for pre-specified group structure. Of all latent factors, ER identifies a specific subset of latent factors that can be used to infer causal associations with the property/outcome of interest. Critically, ER makes no assumptions regarding the underlying data generating mechanisms and can be broadly used across multi-omic datasets (Methods, Supplementary Note 1).

Formally, ER is a latent factor regression in which the unobservable factors Z influence linearly both the response Y and the data X. Its novelty lies in the formulation that enable the latent factors Z to be meaningfully interpreted.

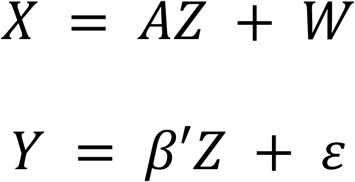

Here, X is the matrix of observables and belongs to a high (*p*) dimensional space (dimensionality of X is *p* x *n,* where n is the number of observations/samples). X is decomposed into the allocation matrix A of dimension *p* x *K* and Z is the latent factor matrix of dimension *K* x *n* i.e., it reduces X from a *p* to a *K* dimensional space (Supplementary Note 1). The matrix Z is used to regress to Y i.e., the regression coefficients correspond to *Z*’s. *W* and *ε* are independent error terms (Supplementary Note 1). This formulation helps cluster the observables (*X’*s) into overlapping clusters/latent factors (Z’s) in a data-dependent fashion and then identify which of the latent factors are significantly associated with, and can be used to infer outcome.

ER comes with two key provable statistical guarantees. The first step is to decompose the matrix of observables X into the latent factor matrix Z. To do this, the membership matrix A needs to be identifiable, up to a *K* × *K* signed permutation matrix. The first guarantee ensures this – we prove that the allocation matrix (A) is indeed identifiable upto a *K* x *K* signed permutation matrix under the assumption that there are at least 2 observables anchoring each latent factor (Supplementary Note 1). This is a reasonable assumption as the model only requires each latent factor to be defined by two observables that are not associated with other latent factors, all other observables may or may not be associated with multiple latent factors. This allows for the identification of group structure from the observables entirely in a data-dependent fashion, without the need to incorporate any prior knowledge. The second guarantee relates to the identifiability of the regression coefficients. We also prove that the coefficient matrix is identifiable up to a signed permutation matrix (Supplementary Note 1), ensuring that the model can rigorously infer significant latent factors driving outcome. Overall, ER is a first-in-class interpretable machine learning framework for uncovering significant latent factors associated with any system-wide property/outcome of interest.

### Composite Regression – a framework to uncover observables driving outcomes from significant latent factors

While the significant latent factors uncovered by ER provide insights into the interplay of the different observables driving outcome, in some contexts their complexity can prove challenging. In these instances, smaller sets of observables underlying outcome are desirable. Currently, regularization is widely used to identify a sparse set of observables (biomarkers). However, regularization-based approaches such as LASSO or Elastic Net uncover predictive biomarkers that may simply be correlative. Given that ER identifies latent factors significantly driving outcome, we sought to develop an approach to identify a sparse set of observables from the significant latent factors identified by ER (Fig. 1c). Using L1-regularization on the significant latent factors identified by ER allows us to identify a sparse set of observables, within these factors, tied to outcome. We term this ER-derivative-approach Composite Regression (CR) (Fig. 1c). As the sparse set of observables delineated by CR are selected from those that lie within the significant latent factors, unlike LASSO-based biomarkers, these are no longer simply predictive, but capture causal relationships that can be used to infer the underlying mechanistic basis of outcome. Together, ER and CR provide a highly prioritized set of significant latent factors and associated observables, which can be used both for inference of underlying cellular/molecular mechanisms as well as corresponding biomarkers.

### ER outperforms state-of-the-art approaches on simulated high-dimensional datasets

We investigated the performance of ER on simulated data (Supplementary Note 1), comparing it with a suite of state-of-the-art approaches including the least absolute selection and shrinkage operator (LASSO) (13), partial least squares (PLS) regression (15), and principal components/factors regression (PFR) (16). We evaluated the performance of ER, PFR, PLS and Lasso change across a range of parameters for original dimensionality (*p*), reduced dimensionality of the dataset (i.e., *K*) and the signal-to-noise ratio (*SNR*). We varied these parameters one at a time and computed the corresponding prediction risk (mean squared error) on data not used to build the model (Supplementary Note 1). We did a grid search on the relevant parameters and found that the prediction error for all four methods deteriorates as *K* increases or the *SNR* decreases (Figs 1d-1f). This indicates that prediction becomes more difficult for large *K* and small *SNR*. On the other hand, ER, PFR and Lasso performed better as *p* increases.

Among the four methods, ER systematically had the smallest prediction error in all settings and PLS had the worst performance in most settings. Furthermore, PFR failed to accurately identify *K* and tended to a very low and sub-optimal *K* in most scenarios (Figs. 1d-1f). This also indicates that, for principal component regression approaches, detecting *K* requires larger SNR i.e., the other approaches are able to accurately detect *K* at lower SNRs. In a moderate SNR regime PFR has comparable performance to ER (Fig. 1d). However, as *K* increases, the advantage of ER becomes considerable, which supports the fact that PFR only has guarantees for fixed K (Fig. 1e). Finally, the performance of PFR is more sensitive to the SNR compared to the other three methods (Fig. 1f). Overall, ER worked very well for very high *p*, was able to accurately identify *K*, and did not have a significant reduction in performance even at lower SNR regimes (Figs. 1d-1f), outperforming state-of-the-art approaches on one or more of these fronts. ER functions counter-intuitively when challenged by the “curse of dimensionality” (i.e., having higher dimensionality is worse as it induces higher variance and can lead to overfitting). The higher dimensionality of the datasets generates more features that provide additional information which are used by ER to predict the latent factors (*Z*) more accurately, thereby overcoming the “curse of dimensionality”.

### Inferring causal factors underlying immunosenescence in a vaccine response

A recent study comprehensively profiled cellular and molecular responses induced by the shingles Zostavax vaccine in a cohort comprising both younger adults and elderly subjects (17). The high dimensional multi-omic analysis included immune cell frequencies and phenotypes, as well as transcriptomic, metabolomic, cytokine and antibody analyses. The vaccine induced robust antigen-specific antibody titers as well as CD4^+^ but not CD8^+^ T cell responses (17). Using a multiscale, multifactorial response network, the authors identified associations between transcriptomic, metabolomic, cellular phenotypic and cytokine datasets which pointed to immune and metabolic correlates of vaccine immunity (17). Interestingly, differences in the quality of the vaccine-induced responses by age were also noted (17). We hypothesized that a method based on latent factors rather than measurables would improve the delineation of components that underlie the quality and magnitude of the vaccine-induced responses. If so, then such a method would be able to leverage the differences in vaccine-induced responses and accurately predict age as the system-wide property of interest. The latent factors identified in this manner could then provide insights into the cellular and molecular basis of age-induced immunosenescence manifested by diminished responses to the Zostavax vaccine.

To explore the above formulation of immunosenescence as a predictor of age, we first applied a suite of state-of-the-art approaches LASSO, PLS and PFR on the entire spectrum of vaccine-induced responses to predict age (Fig. 2a). As most subjects in the cohort were in 2 distinct age groups – adults under 40 and elderly people over 60, we first sought to explore the performance of LASSO, PLS and PFR in predicting the two age groups as binary categorical variables i.e., younger adults and elderly. The predictive performance of all methods was evaluated in a stringent leave-one-out cross-validation (LOOCV) framework (Methods). We have previously demonstrated that on such multi-omic datasets, cross-validation is a gold standard to evaluate model performance with data held out (5, 6, 8). In a LOOCV framework, we found that PFR had no predictive power (AUC < 0.5), while LASSO and PLS had weak predictive power in predicting age as a categorical variable (Fig. 2b, AUCs = 0.63 and 0.60 respectively). The ROC curve for LASSO had an interesting shape. It attained a true positive rate of ~0.4 at a false positive rate of ~0.15, but beyond that it was essentially no better than random (Fig. 2b). This observation is consistent with the observation that differences in an age-associated MMRN were driven by only a subset of elderly vaccinees (17). Thus, a purely predictive modeling approach like LASSO can leverage these relatively straightforward differences to accurately predict age for a subset of the vaccinees, but fails to predict age for others. We then compared these methods to the performance of ER and CR. In a matched, LOOCV framework, ER and CR were very accurate at predicting age (Fig. 2b, AUCs = 0.79 and 0.77 respectively, *P* < 0.01).

**Figure 2.**
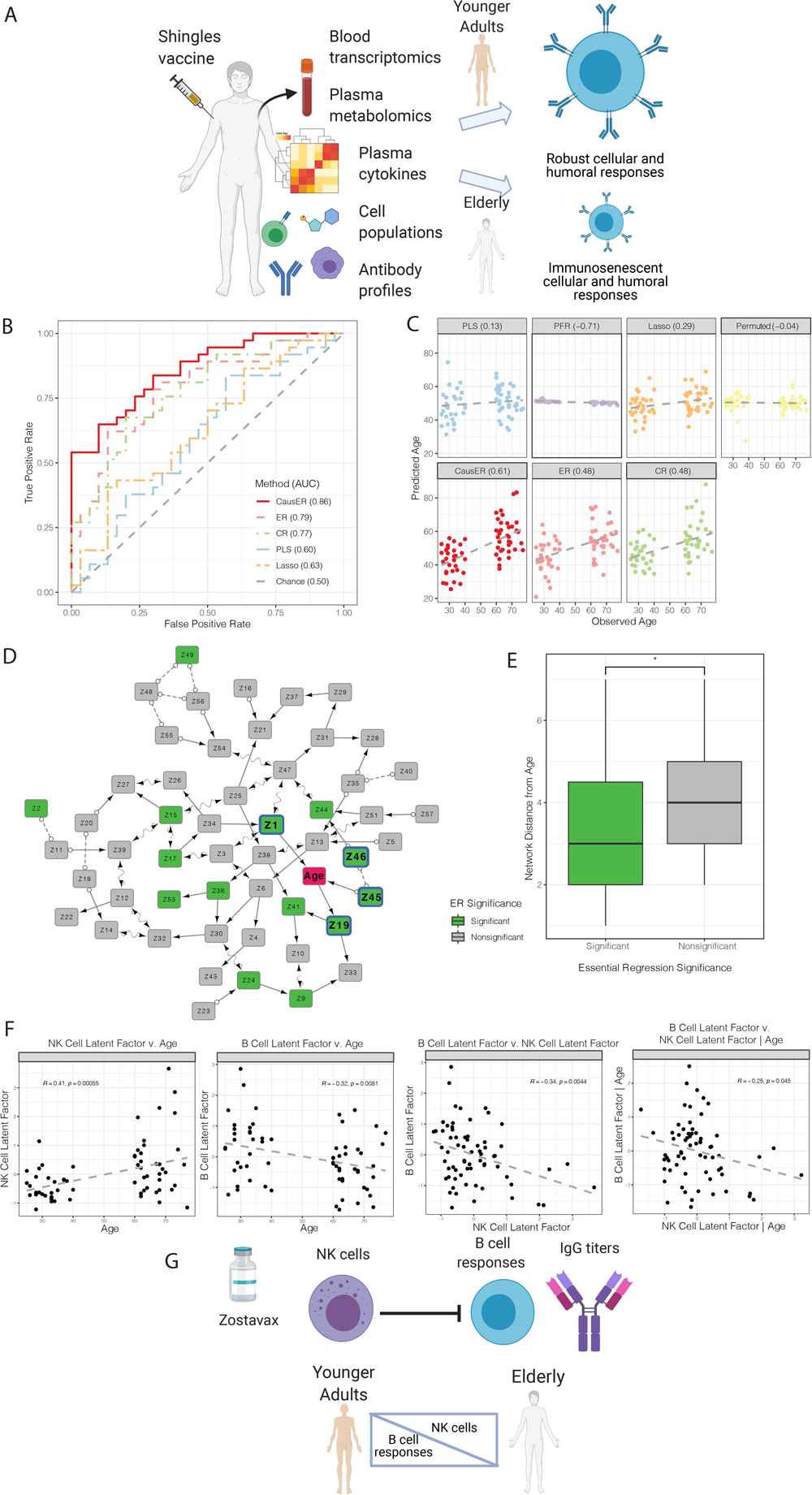
Identifying causal signatures of age-induced immunosenescent responses to the Zostavax vaccine. a) Schematic summarizing the input data and the problem of interest. b) ROC curves for the different methods at discriminating between elderly and younger adults in a LOOCV framework. c) Pearson correlations of the different methods at predicting age as a continuous variable, as measured in a LOOCV cross-validation framework. d) CausalMGM on all Z’s identified by ER. Markov blanket highlighted with a blue border and bolder fonts. A directed edge X --> Y indicates X is a cause of Y, while a bidirected edge X <-> Y indicates the presence of a latent confounder that is a common cause of X and Y. A partially oriented edge X o-> Y indicates that Y is not a cause of X, but either X or a latent confounder causes Y. Unoriented edge indicates directionality couldn’t be inferred for that edge. e) Network distances in the causal graph generated by CausalMGM of the significant and non-significant Z’s identified by ER from the outcome variable of interest. f) Mechanistic insights obtained from ER.

We then coupled ER to causal inference analyses on the ER-identified significant latent factors using directed graphical models(18). Directed Acyclic Graphs (DAGs) are sometimes referred to as Causal Graphs, because under certain assumptions the learned DAGs from observational data (Markov equivalence classes) asymptotically represent the true data-generating causal graph. Although these algorithms have shown considerable success in analyzing many biological processes and biomedical problems (19–23), including biomarker selection and classification (24–26), scalability limits the datasets that they can be applied (27, 28). Here, we use the causal learning algorithm for mixed data, CausalMGM,(19, 29) only on the significant latent factors delineated by ER, to overcome the scale limitation. By applying CausalMGM only on the significant latent factors, we greatly reduce the dimensionality of the input dataset while preserving the information of individual (correlated) variables in the latent factors. Thus, CausER (<u>Caus</u>alMGM on the significant latent factors from <u>ER</u>) prioritizes further within the significant latent factors (Methods, Supplementary Note 1) by virtue of their direct connections to the outcome in the graphical model. Furthermore, it predicts potential cause-effect relationships between the latent factors and the property/outcome of interest, which leads to hypotheses generation. While CausER was the best predictor of age as a categorical variable (AUC = 0.86, *P* < 0.01). Together, these results demonstrate that while LASSO, PLS and PFR fail to accurately predict age from Zostavax-induced vaccine responses, ER, CR and CausER can overcome this challenging problem by leveraging non-trivial differences in latent factors comprised of discrete sets of measurables.

Next, we evaluated whether these methods could predict actual age as a continuous variable beyond the categorical classifiers of younger adults and the elderly. As before, performance was measured in a rigorous cross-validation framework (Methods). Using the vaccine-induced responses, PFR was not at all predictive of age (Fig. 2c, Pearson r = −0.71; Fig. S2, Spearman r = −0.82). LASSO and PLS had poor performance in predicting age as a continuous variable (Fig. 2c, Pearson r = 0.29 and 0.13 respectively; Fig. S2, Spearman r = 0.25 and 0.09 respectively). In fact, the predictive power of PLS and PFR were not significantly different from a negative control model built on permuted data (Fig. 2c). However, both ER and CR were significantly predictive of age as a continuous variable (Pearson *r* = 0.48 for both, Spearman r = 0.44 and 0.49 respectively, *P* < 0.01 Fig. 2c, Fig. S2), and as in the previous instance, CausER had the best performance in predicting age as a continuous variable (Pearson *r* = 0.61, Spearman r = 0.59, *P* < 0.01 Fig. 2c, Fig. S2). Together, these results demonstrate that while state-of-the-art methods including LASSO, PLS and PFR fail to predict age either as a categorical or a continuous variable, all three of the new approaches that are based on latent factors – ER, CR and CausER, are able to do so reasonably accurately based on the multi-omic profiles of vaccine-induced responses.

We next explored the likely causal relationships among the latent factors that lead to age-induced immunosenescence and diminished responses to the Zostavax vaccine. CausalMGM was used to construct a causal graph with all latent factors identified in the latent model identification step of ER (Fig. 2d). Notably, majority of the significant latent factors identified by ER were seen to proximal to the outcome variable (age) in the causal graph. Importantly, all 4 latent factors in the Markov blanket generated by CausalMGM were also identified as significant by ER (Fig. 2d). Overall, the significant latent factors revealed by ER had significantly lower network distances (i.e., had stronger cause-effect relationships) from age compared to the non-significant latent factors (Fig. 2e, *P* < 0.05). These results demonstrate that the cause-effect relationships identified by ER are validated by CausalMGM. Importantly, while CausER hits are identified via the sequential application of ER and CausalMGM respectively, the order is critical with ER being the key first step. Without the two-stage dimensionality reduction (first from observables to latent factors, and then the identification of significant latent factors) afforded by ER, running CausalMGM or other allied causal graphical models on the initial set of observables would be computationally intractable.

The prioritized CausER hits (Fig. 2d) i.e, significant latent factors identified by ER that are also in the Markov blanket of the outcome variable (age) in the causal graph generated by CausalMGM comprised antigen-specific IgG titers (Z1), a metabolic module (Z19), B cell (Z46) and NK cell frequencies (Z45). CausER provides both prioritized cause-effect relationships and directions of these relationships. While the latter relates to mathematical conditional independence relationships (Methods), the former provides prioritized mechanistic insights. While the lowering of titers with age is expected and has been previously reported(17), CausER revealed a likely cause-effect relationship between altered B cell and NK cell numbers and immunosenescence. To further dissect the nature of this relationship, we examined correlations between NK cells, B cells and age. We found that NK cells significantly increased, while the numbers of B cells significantly decreased with age (Fig. 2f). More interestingly, there was a significant negative correlation between NK cells and B cells (Fig. 2f), and the correlation remained significant even after correcting for age (Fig. 2f). Our results suggest a novel basis of human immunosenescence in the context of vaccine responses (Fig. 2g). This could involve a previously described mechanistic linkage between NK cells and a weaker germinal center (GC) response in a murine model (30). NK cells can inhibit CD4 T cell responses including those of T follicular helper cells in a perforin-dependent manner; this leads to a weaker GC response diminished antibody titers and affinity maturation (30, 31).

### Uncovering latent factors that distinguish immune system states of term and pre-term infants

Next, we focused on a multi-omic longitudinal cohort that analyzed immune cell populations and plasma proteins in 100 newborn children during their first 3 months of life (32) (Fig. 3a). Striking differences were observed in immune parameters between preterm and term children at birth. However, the immune trajectories appeared to achieve a stereotypic convergence within the first 3 months of life (32) (Fig. 3a). We hypothesized that ER might be able to uncover latent factors that distinguish immune system states of term and pre-term infants after 3 months of life and therefore reveal features that could impact later life (Fig. 3a). As expected, based on the striking differences at birth between term and pre-term children, all methods (LASSO, PLS, PFR, ER, CR and CausER) were be able to discriminate between these 2 groups using immune parameters measured in the first week of life (Fig. S4). All model performances were measured in a rigorous cross-validation framework (Methods). However, given the stereotypic convergence in the first 3 months (12 weeks) of life (32), we found that PLS and PFR were unable to accurately discriminate between term and pre-term children using immune parameters measured at 12 weeks of life (Figs. 3b-3c). However, LASSO was able to accurately distinguish between term and pre-term births using the 12-week profiles (Figs. 3b-3c), suggesting that despite broad convergence, a small subset of immune parameters still remain different term and pre-term infants between at 3 months of life. More importantly, ER and CR were able to accurately discriminate between term and pre-term births using immune profiles at 3 months of life, significantly better than other methods (Figs. 3b-3c, P < 0.01). ER identified only 2 significant latent factors, and based on CausalMGM analyses, one of these 2 significant latent factors was in the Markov blanket i.e., for this dataset, this single latent factor was the sole CausER hit (Fig. 3d).

**Figure 3.**
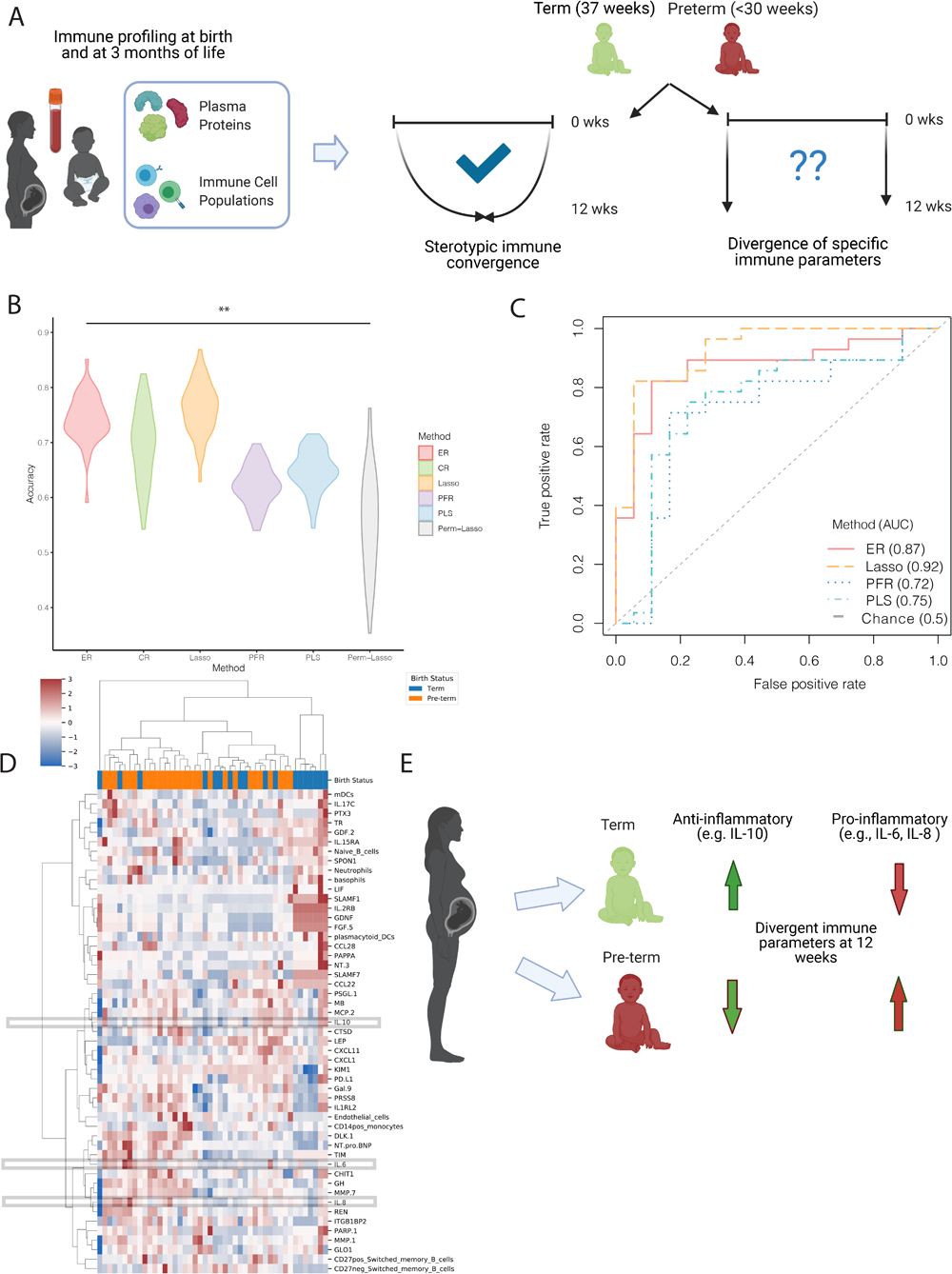
Uncovering specific immune parameters from term and pre-term infants that do not achieve stereotypic convergence. a) Schematic summarizing the input data and the problem of interest. b) Classification accuracy of the different methods at discriminating between term and pre-term births using immune profiles at 3 months after birth, measured in a replicated *k*-fold cross validation framework. c) ROC curves for the different methods at discriminating between term and pre-term births as measured in a LOOCV framework. d) Heatmap of features (plasma proteins and immune cells) in the single hit (significant Z identified by ER in the Markov blanket of outcome). e) Mechanistic insights obtained from ER.

We visualized the immune cell populations and plasma proteins in this hit (Fig. 3d). These profiles had clearly remained divergent even at 3 months of life (Fig. 3d) despite the broad stereotypic convergence of most other immune parameters. At 3 months of life, term infants had an anti-inflammatory milieu including high IL-10, while pre-term infants had a pro-inflammatory milieu including elevated IL-6 and IL-8 (Fig. 3d). These findings agree with a previous study that IL-10 is highly expressed in the uterus and placenta and has a key role in controlling inflammation-induced pre-term labor in a murine model (33). Furthermore, regulatory B cells are a key source of IL-10 and appear to be important in sustaining pregnancy till term (34–36). It is also known that modulation of pro-vs-anti-inflammatory environments by relevant cytokines and chemokines at the maternal-fetal interface (decidua) is a critical component of the bifurcation between term and pre-term births (34). Thus, our analyses of immune system states of term and pre-term infants at 3 months of life revealed that pre-term infants had a pro-inflammatory state, while term infants had an anti-inflammatory state (Fig. 3e). These findings could have long-term implications for the health of pre-term infants.

## Discussion

Over the last two decades, while there have been rapid advances in high-throughput experimental technologies to generate deep molecular profiles, computational analyses of these high-dimensional datasets have primarily focused on biomarker discovery (37). This is because rigorous statistical approaches for analyzing high-dimensional datasets, such as regularized regression and bootstrap aggregated classification, are focused on uncovering predictive biomarkers which may simply be correlative surrogates of outcome or system-wide property but unrelated to the underlying causal factors. Incorrect extrapolation of insights derived from biomarker-based approaches can lead to perturbation experiments with low success. Alternatively, efforts to move beyond biomarkers to mechanistic insights often use biological priors, which may be incomplete or suffer from sampling/study biases (38). Further, while there have been advances in causal modeling (39), existing approaches are difficult to apply to high-dimensional datasets due to the computational intractability of applying these approaches on (18) and the multi-collinearity of the data. The methods presented in this manuscript address this fundamental limitation in systems biology. ER is a first-in-class machine learning method that can both handle high-dimensional multi-omic datasets with co-linear variables and prioritize cause-effect relationships between the input features and the outcome of interest. Our framework is also complementary to modern approaches that combine multi-omic datasets with prior knowledge to uncover causal relationships (40). ER generates mechanistic hypotheses solely based on latent factors identified from multi-omic data without the incorporation of any prior knowledge. It is thus applicable in contexts where prior knowledge is weak or unavailable, and is not limited by the nature and quality of available prior knowledge.

The ER framework pushes the envelope on multiple key challenges in systems biology. First, it establishes a rigorous framework with provable statistical guarantees that explores a large space of higher-order relationships from high-dimensional features and uncovers latent factors tied to the outcome variable via directed cause-effect relationships. Second, unlike existing causal reasoning approaches that are constrained by the size of the input data, ER can be applied to modern high-dimensional datasets. The time complexities of the different steps are essentially quadratic and not exponential like some other causal reasoning approaches. Third, ER makes no assumptions regarding data-generating mechanisms and ER can integrate multi-omic datasets to capture the interplay across a plethora of biological processes at multiple scales of organization of the system.

An important elaboration of our framework is the sequential use of two orthogonal methods for statistical inference, ER and causal graphical modeling. These methods have different theoretical bases and assumptions and yet the ER hits are validated by CausalMGM, underscoring the robustness of our approach. The order is critical with ER being the key first step offering a two-stage dimensionality reduction – first from observables to latent factors, and then the identification of significant latent factors. Without these two steps, the application of causal graphical models on the initial set of observables would be computationally intractable due to the high dimensionality of the dataset. Thus, ER solves a long-standing limitation with causal graphical modeling approaches, and enables for the first time, causal inference on high-dimensional data.

Here, we applied ER to diverse contexts. First, we applied ER on simulated datasets and demonstrated that it performed better than LASSO, PLS, PFR across a range of parameter settings. Next, we utilized ER on two recent human systems immunology studies that had generated high-dimensional multi-omic profiles. Using ER, we were able to address key questions that had not been the focus of the original studies, in part because of limitations of methods used. Such questions could now be addressed by the methodological advances of ER over state-of-the-art approaches. We demonstrated that ER significantly outperforms PFR and PLS across contexts, and either outperform or match LASSO in terms of predictive performance. While we used three examples to illustrate the superior performance of ER, these methods come with broad theoretical guarantees to outperform PLS, PFR and LASSO across contexts (Methods, Fig. S5). Further, while the existing methods simply identify correlates, ER provides mechanistic insights, some of which are consistent with prior knowledge, while others are novel. ER can also be used for noisier and older datasets not generated using state-of-the-art methods (Supplementary Note 2). Our findings have broad implications across domains in systems biology and are likely to transform both computational workflows used to analyze multi-omic datasets and downstream experiments designed based on the insights gleaned via these analyses.

## Methods

We provide brief descriptions of the methods, associated tuning parameters, cross-validation strategies and data pre-processing in this section. Additional details are included in Supplementary Note 1.

### Processing of human systems immunology datasets

For the dataset of multi-omic responses to the Zostavax vaccine, we included the following multi-scale measurements of immune state: IgG titers, blood transcriptional modules, metabolic clusters, CD4+ T cell populations, TFH cell populations, flow cytometry cell populations, cytokine profiles, and IFN T cells. We used subject age as the response variable for *n* = 72 subjects. We excluded features that have missing values for more than a half of subjects. We also exclude 5 subjects that have no observed features. The remaining data sets are merged via the unique ids of subjects. The final data set contains *p* = 1721 features of *n* = 67 subjects.

For the dataset of term and pre-term infants, we included all available immune parameters as features and only removed clinical metadata (such as. “gender”, “mode of delivery”, “family” etc. The final data set we use has *n* = 183 samples and *p* = 282 features with 56 samples from week 1 and 46 samples from week = 12. The response is binary, either “Control” (representing term) or “Premature” (representing pre-term). We use the 5-NN to impute the missing values.

### Cross-validation

Two cross-validation techniques were used to assess the predictive performance of the different methods – 1) replicated 10-fold cross-validation, and 2) leave-one-out cross-validation. 1) To assess the accuracy of the classifiers for the term / pre-term immune profile, 50 replicates of nested 10-fold cross-validation were performed. For each replicate, we independently ran each of the methods and assessed the predictive accuracy. For ER, the latent factors were learned on each fold and each replicate, and the regression and final latent factor selection were repeated. For CausER, a causal model was learned over the latent factors selected as significant by ER for each fold and replicate. The average cross-validation accuracy across the 10 folds was calculated for each of the 50 replicates. 2) For all three datasets, we also performed leave-one-out cross-validation to assess the accuracy of each method. In leave-one-out cross-validation, each sample in the dataset is held out as the predictive models are trained on the remaining samples, and then the held-out sample is predicted with the trained models. Assessment of model performance (AUC) was done with the set of predictions for the left-out values.

### ER

The first step in ER is the estimation of all latent factors. The identification of latent factors is unsupervised. This is done based on the empirical sample covariance matrix using a three-step procedure. The first step involves the identification of latent variable structure using the sample covariance matrix. A key component of this step is the identification of *K* (reduced dimensionality) from *p* (original dimensionality). The second step involves inference of the clusters – each cluster (latent factor) is anchored by at least 2 pure variables. Variables that are associated with multiple clusters are designated mixed variables. The third involves determination of the overall allocation matrix based on the cluster assignments in the earlier step. Formal descriptions of all 3 steps are provided in Supplementary Note 1, Section 2. After the identification of Z’s, betas linking the Z’s to Y are estimated using a theoretical framework we recently established for estimation in latent factor regression models(41). This is the supervised part of the ER algorithm. A detailed description of the estimation procedure is provided in Supplementary Note 1, Section 2.

### CR

CR utilizes a 2-step procedure. First, it uses ER to identify significant Z’s as described above. Then, it uses LASSO on the X’s associated only with the significant Z’s to identify a sparse basis for the system-wide property/outcome of interest. For LASSO on the significant Z’s identified by ER, lambda is tuned using k-fold cross-validation. The lambda tuning is specific to a given fold for a given replicate and utilizes only the fold-specific training data.

### ER coupled to CausalMGM

We implemented CausalMGM as previously described(18) on all Z’s for the Zostavax dataset and only the significant Z’s identified by ER for the term/pre-term dataset. Briefly, when constructing the causal model, we first learned an undirected graphical model with MGM(42)/GLASSO(43). The optimal regularization parameters were selected based on graph stability using StEPS(29)/StARS(44). The resulting undirected graph was then used as an initial graph for performing causal inference with the FCI algorithm. To build a predictor of the outcome variable, the Markov blanket was used. The Markov blanket was defined as the set of variables that, when conditioned on, make the response variable independent of every other variable in the dataset according to the structure of the causal graph. For a DAG, this comprises the parents, children, and spouses (other parents of the children) of the response variable.

### Implementation of LASSO, PLS and PFR

LASSO was implemented using glmnet in R with parameter tuning done in a manner analogous to that described above for ER and CR. If no feature was selected by LASSO in a specific fold for a given replicate, we randomly selected 5 features (only for that fold in that replicate) and use an ordinary least squares estimator. Thus, the feature selection in each case is specific to each fold for a given replicate; this is the most stringent and unbiased way to evaluate model performance. PLS was implemented using the plsr function in R with the number of components selected by the default function selectNcomp. For PFR, which regresses *Y* on the first *K* principal components of *X*, the number of principal components *K* is selected based on the ratios of non-decreasing eigenvalues of *X*’\**X*/*n*, using previously established criteria(45).

## Code Availability

Detailed code and documentation for ER and CR and CausER are available at https://github.com/bingx1990/Application-of-ER-and-CausalMGM.git

## Author Contributions

J.D. designed the study, and oversaw all aspects of it. X.B., F.B. and M.W. jointly conceived the theoretical basis of the ER framework. J.D. and X.B. jointly conceived the application of the ER and CR frameworks to real data. J.D., P.B. and H.S. jointly conceived the CausER framework. X.B. and T.L. implemented the ER, CR and CausER frameworks, and carried out all computational analyses. J.D. and H.S. interpreted the results. J.D., H.S. and P.B. wrote the main text. X.B., T.L., F.B. and M.W. wrote the supplementary methods including formal proofs.

## Supplementary Figure Legends

**Fig. S1 – Spearman correlations of the different methods at predicting age as a continuous variable, as measured in a LOOCV cross-validation framework.**

**Fig. S2 – Performance of ER in discriminating between term and pre-term births using immune profiles at week 1 after birth**

a) Classification accuracy of the different methods at discriminating between term and pre-term births using immune profiles at 1 week after birth, measured in a replicated k-fold cross validation framework

b) ROC curves for the different methods at discriminating between term and pre-term births as measured in a LOOCV framework.

**Figure S3 – Identifying differences in vaccine-induced transcriptomic profiles over time**

a) Schematic summarizing the input data and the problem of interest.

b) Ternary classification accuracy of the different methods at discriminating among G1, G2 and G3 in a replicated *k*-fold cross-validation framework.

c) Confusion matrix summarizing the performance of the different methods at discriminating among G1, G2 and G3 in a LOOCV framework.

d) ROC curves for the different methods at discriminating between G3 and (G1 & G2) combined in a LOOCV framework.

e) ROC curves for the different methods at discriminating between G3 and G1 in a LOOCV framework.

f) Fraction of true G3 correctly classified as G3 (as measured in a LOOCV framework).

g) CausER graph – CausalMGM on the significant Z’s from ER. Markov blanket highlighted with a blue border and bolder fonts. A directed edge X --> Y indicates X is a cause of Y, while a bidirected edge X <-> Y indicates the presence of a latent confounder that is a common cause of X and Y. A partially oriented edge X o-> Y indicates that Y is not a cause of X, but either X or a latent confounder causes Y. Unoriented edge indicates directionality couldn’t be inferred for that edge.

h) Heatmap of genes in CausER hits (significant Z’s in the Markov blanket) for G1 and G3 samples.

i) Heatmap of genes in CausER hits (significant Z’s in the Markov blanket) for G1, G2 and G3 samples

**Supplementary Note 1**

Detailed descriptions of the theoretical underpinnings, associated proofs of ER, CR and CausER. The file also describes details of the applications of ER, CR, CausER, LASSO, PLS and PFR to the different datasets of interest.

**Supplementary Note 2**

Applying ER to datasets generated using older technologies

