## Supplementary figures and images for "Essential Regression - a generalizable framework for inferring causal latent factors from multi-omic human datasets"

### Supplementary Figures

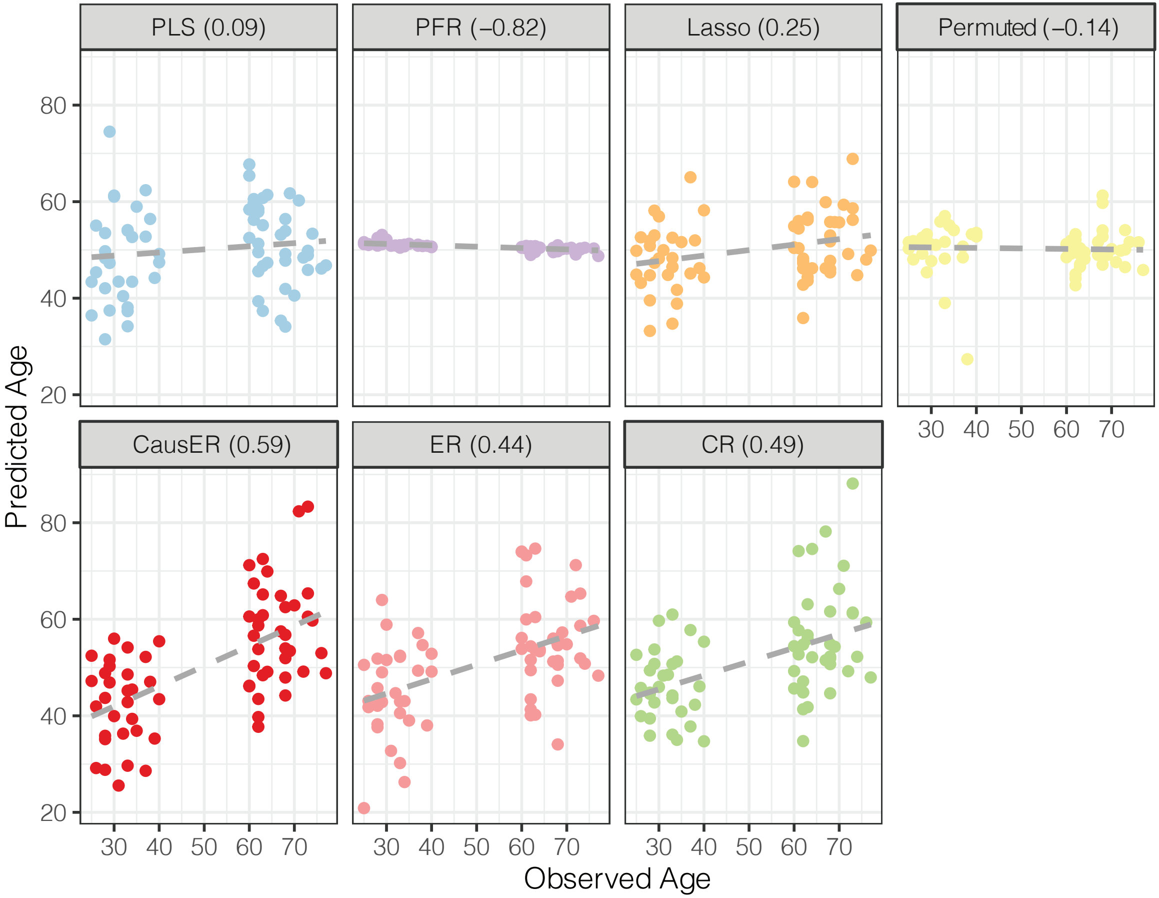

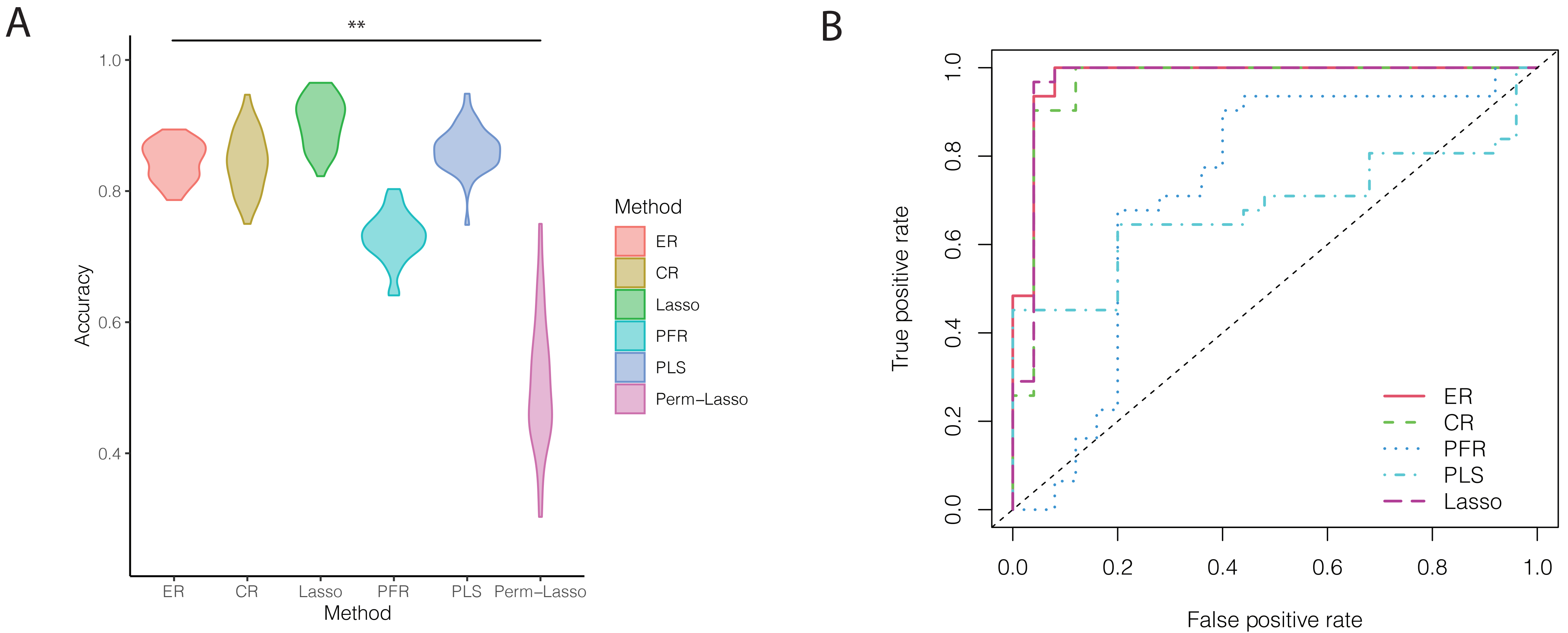

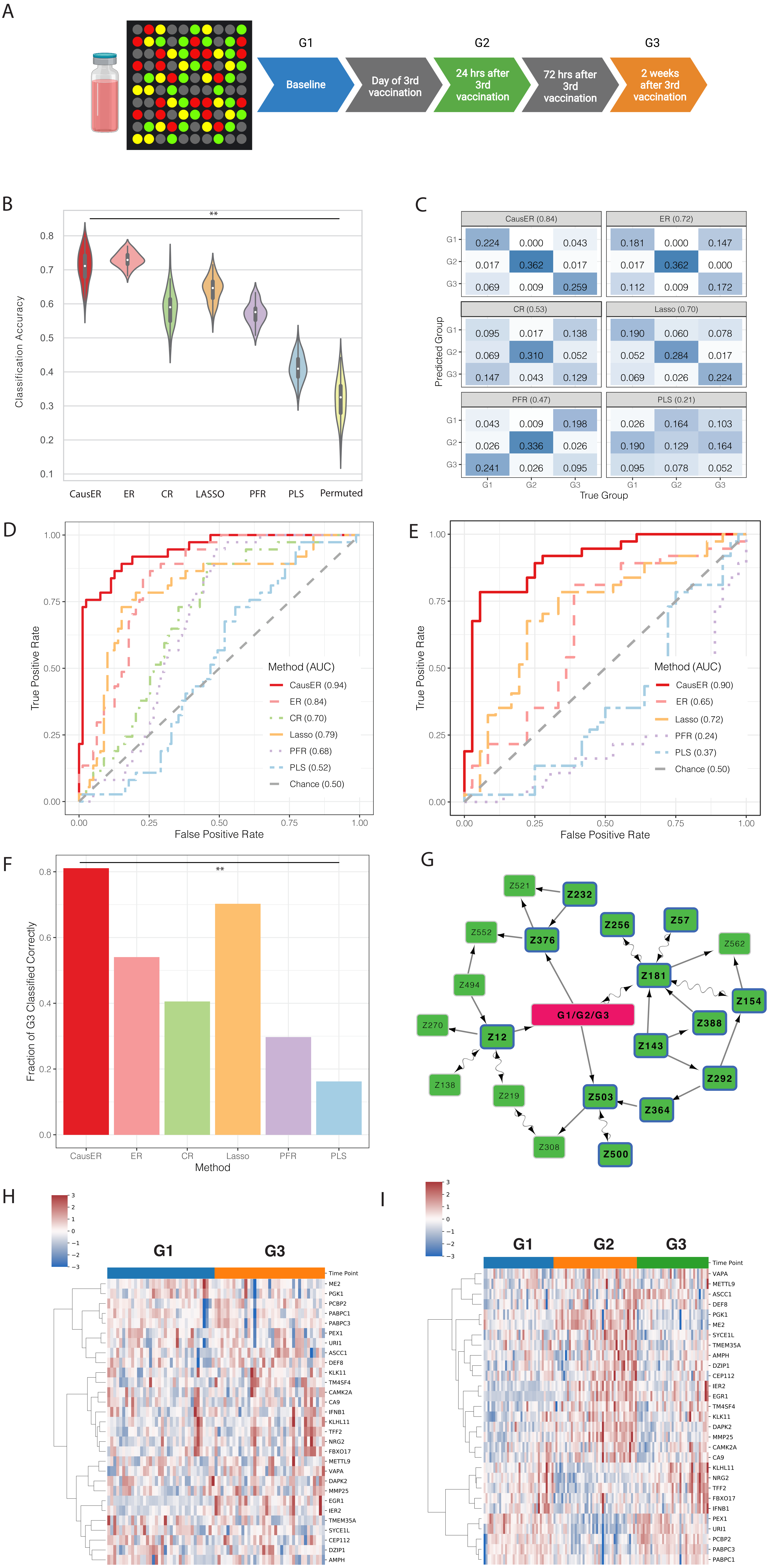
