## Supplementary Note 1 for "Essential Regression - a generalizable framework for inferring causal latent factors from multi-omic human datasets"

#### 1 Latent factor model formulation

We assume that the data $(\mathbf{X}, \mathbf{Y}) \in (\mathbb{R}^{n\times p}, \mathbb{R}^n)$ are i.i.d. realizations of the random vector $(X, Y) \in (\mathbb{R}^p, \mathbb{R})$ which follows the factor regression model

$$Y = \beta^\top Z + \varepsilon \tag{1.1}$$

$$X = AZ + W. \tag{1.2}$$

The latent, unobserved, random vector $Z \in \mathbb{R}^K$, with $K < p$, is independent of the mean-zero random errors $\varepsilon \in \mathbb{R}$ and $W \in \mathbb{R}^p$. Independence between $\varepsilon$ and $W$ is assumed as well. The $p \times K$ matrix $A$ and the $K$-dimensional vector $\beta$ are deterministic quantities. We let $\mathbb{E}[\varepsilon^2] = \sigma^2$ and assume both $\mathbb{E}[ZZ^\top] = \Sigma_Z$ and $A$ have rank $K$.

Since one of our main interests is to cluster the feature $X$ based on its association with the latent factor $Z$, the membership matrix $A$ needs to be identifiable, up to a $K \times K$ signed permutation matrix. For this reason, we resort to the following conditions on $A$, $\Sigma_Z$ and $\Gamma := \mathbb{E}[WW^\top]$ (1).

**Assumption 1.**

*(A0)* $\|A_{j\cdot}\|_1 \le 1$ *for all* $j \in [p] := \{1, 2, \ldots, p\}$.

*(A1) For every* $k \in [K]$, *there exists at least two* $j \neq \ell \in [p]$, *such that* $|A_{j\cdot}| = |A_{\ell\cdot}| = e_k$.

*(A2) There exists a constant* $\nu > 0$ *such that*

$$\min_{1\le a<b\le K} \left( [\Sigma_Z]_{aa} \wedge [\Sigma_Z]_{bb} - |[\Sigma_Z]_{ab}| \right) > \nu.$$

*(A3) The covariance* $\Gamma := \mathbb{E}[WW^\top]$ *is diagonal with bounded diagonal entries.*

Model (1.1) – (1.2) together with Assumption 1 is termed as the Essential Regression (ER). Within the ER framework, (1, Theorem 2) and (2, Proposition 1) establish that the model is identifiable. In particular, both $A$ and $\beta$ are identifiable. For the reader's convenience, we restate the results in Appendix A. The uniqueness of $A$ is used to form unique clusters of $X$ based on their associations with $Z$ via

$$G_k := \{j \in [p] : A_{jk} \neq 0\}, \quad \forall k \in [K].$$

On the other hand, the coefficient $\beta$ is used to select significant $Z$ for predicting $Y$

To establish the identifiability results, the key condition is *(A1)* which assumes the existence of at least two features $X$ that are *solely* associated with each latent factor $Z$. These features are termed as *non-mixed variables* and are collected in the set $I \subseteq [p] := \{1, 2, \ldots, p\}$. We further let $\mathcal{I} = \{I_1, \ldots, I_K\}$ denote its partition. Mathematically, we have

$$I_k := \{i \in [p] : |A_{ik}| = 1, A_{ik'} = 0, \text{ for all } k' \neq k\}, \quad \text{for } 1 \le k \le K.$$

The existence of non-mixed variables also renders the latent factors $Z$ interpretable as the meaning of each $Z_k$ can be read off from the corresponding non-mixed features $\{X_i\}_{i\in I_k}$.

Under the Essential Regression, our goals are two-fold:

(a) estimate $\beta$ and select predictive latent factors $Z$;

(b) predicting $Y$.

In the next section, we will describe our approaches for each goal separately.

### 2 Methodology under Essential Regression

#### 2.1 Estimation of $\beta$ and selection of predictive $Z$

##### 2.1.1 Estimation of $\beta$

Let $\widehat{\Sigma} := n^{-1}\mathbf{X}^\top\mathbf{X}$ denote the empirical sample covariance matrix of $X$. Our procedure for estimating $\beta$ is the following.

(1) Obtain estimates $\widehat{K}$, $\{\widehat{I}_1, \cdots, \widehat{I}_{\widehat{K}}\}$, $\widehat{A}_{\widehat{I}\cdot}$ and $\widehat{\Sigma}_Z$ from $\widehat{\Sigma}$ with tuning parameter $\delta$ by using Algorithm 1 in (1). For the reader's convenience, the procedure is stated in Appendix B.

(2) Estimate $\Gamma_{\cdot I}$ by $\widehat{\Gamma}_{\cdot \widehat{I}}$ with

$$\widehat{\Gamma}_{ii} = \widehat{\Sigma}_{ii} - \widehat{A}_{i\cdot}^\top \widehat{\Sigma}_Z \widehat{A}_{i\cdot}, \quad \forall\, i \in \widehat{I}, \qquad \widehat{\Gamma}_{ji} = 0, \quad \forall\, j \neq i. \tag{2.1}$$

(3) Compute

$$\widehat{\Theta} = \left(\widehat{\Sigma}_{\cdot\widehat{I}} - \widehat{\Gamma}_{\cdot\widehat{I}}\right) \widehat{A}_{\widehat{I}\cdot} \left(\widehat{A}_{\widehat{I}\cdot}^\top \widehat{A}_{\widehat{I}\cdot}\right)^{-1}. \tag{2.2}$$

If $\widehat{\Theta}^\top\widehat{\Theta}$ is non-singular, estimate $\beta$ by

$$\widehat{\beta} = \left(\widehat{\Theta}^\top\widehat{\Theta}\right)^{-1} \widehat{\Theta}^\top \frac{1}{n}\mathbf{X}^\top\mathbf{Y}. \tag{2.3}$$

Otherwise, compute

$$\widehat{h} = \frac{1}{n}\left(\widehat{A}_{\widehat{I}}^\top \widehat{A}_{\widehat{I}}\right)^{-1} \widehat{A}_{\widehat{I}}^\top \mathbf{X}_{\widehat{I}}^\top \mathbf{Y} \tag{2.4}$$

and estimate $\beta$ by

$$\widehat{\beta}_d = \arg\min_{\beta \in \mathbb{R}^K} \left\{ \|\beta\|_1 : \|\widehat{\Sigma}_Z\beta - \widehat{h}\|_\infty \leq \mu_1 + \mu_2\|\beta\|_1 \right\} \tag{2.5}$$

for some parameters $\mu_1$ and $\mu_2$.

Algorithm 1 in step (1) requires to choose the tuning parameter $\delta$. Since theoretical order of $\delta$ is $\sqrt{\log(p \vee n)/n}$ under the sub-Gaussian assumption of $Z$, $\varepsilon$ and $W$, we set $\delta = c\sqrt{\log(p \vee n)/n}$ with the leading constant $c$ chosen via the criterion in Section 5.1.1 of (1).

For $\widehat{\beta}_d$, the procedure requires additional tuning parmeters: $\mu_1$ and $\mu_2$. They are all of theoretical order $\sqrt{\log(p \vee n)/n}$. Our extensive simulation suggests to choose $\mu_1 = \mu_2 = 0.5\sqrt{\log(p \vee n)/n}$. Alternatively, they can be chosen via cross-validation by minimizing the loss

$$L(\beta) := \beta^T \widehat{\Sigma}_Z \beta - 2\beta^\top \widehat{h}.$$

##### 2.1.2 Selection of predictive $Z$

When our estimator of $\beta$ is $\widehat{\beta}$, (2, Theorem 4 and Proposition 5) provides the asymptotic distribution of $\widehat{\beta}_k$ for all $1 \leq k \leq K$ with consistent estimates of the asymptotic variances. We thus can construct confidence intervals (CIs) for each $\beta_k$, $1 \leq k \leq K$ and the obtained CIs could be used to select the predictive latent factors $Z$.

When our estimator of $\beta$ is the Dantzig-type estimator $\widehat{\beta}_d$, as mentioned in (2, Remark 3 of version 1), $\widehat{\beta}_d$ adapts to the unknown sparsity of $\beta$. We propose to directly use the support of $\widehat{\beta}_d$ to select predictive $Z$.

#### 2.2 Prediction

We have two procedures for predicting $Y$, described separately in the following two sections.

##### 2.2.1 Essential regression predictor

For predicting $Y$, we adopt the procedure in (3, Section 4.2). Specifically, let $\widehat{\Theta}$ be constructed from (2.2) and compute

$$\widehat{\theta}_{ER} := (\widehat{\Theta}^\top \mathbf{X}^\top \mathbf{X} \widehat{\Theta})^- \widehat{\Theta}^\top \mathbf{X}^\top \mathbf{Y}$$

where $M^-$ denotes the Moore-Penrose inverse of any matrix $M$. For any new data point $(X^*, Y^*)$, we predict $Y^*$ by

$$\widehat{Y}^*_{ER} = \widehat{\theta}_{ER}{}^\top X^*. \tag{2.6}$$

This predictor is termed as ER in our result.

##### 2.2.2 Composite regression predictor

Within the Essential Regression framework, although the significant latent factors $Z$ could be selected, the ER predictor in (2.6) still uses all the features $X$ to predict $Y$. We thus propose a new predictor, called Composite Regression (CR), which uses only the features $X$ that are related with the selected significant $Z$.

Specifically, CR has two steps. In the first step we select the significant factors $Z$ and let $L \subseteq [K]$ be the index set of the selected $Z$. In the second step, we first find the subset of features $X$ that are related with $Z_L$, that is, the set

$$\bar{S} := \left\{ j \in [p] : \|\widehat{A}_{jL}\|_2 \neq 0 \right\}$$

where $\widehat{A}$ is estimated from (B.4) in Appendix B, the procedure proposed in (1). We then regress $\mathbf{Y}$ onto $\mathbf{X}_{\bar{S}}$ via the Lasso approach to obtain the estimated linear coefficient vector

$$\widehat{\theta}_{CR} := \arg\min_{\theta} \|\mathbf{Y} - \mathbf{X}_{\bar{S}}\theta\|_2^2 + \lambda \|\theta\|_1.$$

The estimate $\widehat{\theta}_{CR}$ could be used to select predictive features associated with those significant factors $Z$. Furthermore, we propose

$$\widehat{Y}^*_{CR} := \widehat{\theta}_{CR}^\top X^* \tag{2.7}$$

to predict $Y^*$.

It is worth mentioning that the difference between CR and Lasso is that CR regresses $\mathbf{Y}$ onto $\mathbf{X}_{\bar{S}}$ based on the selected significant $Z_L$ whereas Lasso regresses $\mathbf{Y}$ onto *all* the features $\mathbf{X}$. Hence the selected features $X$ from CR are associated with the predictive factors, a desirable property that Lasso does not enjoy.

#### 2.3 Prediction with Essential Regression on synthetic data

In this section[1] we generate synthetic data to compare the prediction performance of ER relative to PFR, PLS and the Lasso.

[1] This section is modified based on Section 5.2 in (4)

##### 2.3.1 Data generating mechanism

We start with the description of our data generating mechanism. We first describe how we generate $A$, $\Sigma_Z$, $\Gamma$, and $\beta$. Recall that $A$ can be partitioned into $A_I$ and $A_J$.

To generate $A_I$, we set $|I_k| = m$ for each $k \in [K]$ and choose $A_I = \boldsymbol{I}_K \otimes \mathbf{1}_m$, where $\otimes$ denotes the kronecker product. Each row $A_{j\cdot}$ of $A_J$ is generated by first randomly selecting its support with cardinality $s_j$ drawn from $\{2, 3 \ldots, \lfloor K/2 \rfloor\}$ and then by sampling its non-zero entries from $N_{s_j}(0, D)$. The matrix $D$ satisfies $\text{diag}(D) = (1/s_j, \ldots, 1/s_j)$ and $D_{ab} = \zeta^{|i-j|}/s_j$ for any $a \neq b$ with given parameter $\zeta \in [0, 1]$. In the end, we rescale $A_J$ such that the $\ell_1$ norm of each row is no greater than 1.

To generate $\Sigma_Z$, we set $\text{diag}(\Sigma_Z)$ to a $K$-length sequence from 2.5 to 3 with equal increments. The off-diagonal elements of $\Sigma_Z$ are then chosen as $[\Sigma_Z]_{ij} = (-1)^{(i+j)}([\Sigma_Z]_{ii} \wedge [\Sigma_Z]_{jj})(0.3)^{|i-j|}$ for any $i \neq j \in [K]$. Finally, $\Gamma$ is chosen by randomly sampling its diagonal elements from $\text{Unif}(3, 5)$ and the entries of $\beta$ are sampled independently from Unif(0, 1).

We generate the $n \times K$ matrix $Z$ and the $n \times p$ noise matrix $W$ whose rows are i.i.d. from $N_K(0, \Sigma_Z)$ and $N_p(0, \Gamma)$, respectively. Then we set $X = ZA^T + W$ and $Y = Z\beta + \varepsilon$ where the $n$ components of $\varepsilon$ are i.i.d. $N(0, 1)$. For each setting, we repeat generating $(X, Y)$ 50 times and record the corresponding results.

Below we investigate how the prediction errors of ER, PFR, PLS and Lasso change as we vary $p$, $K$ and the signal-to-noise ratio (SNR) one at a time. The performance metric is based on the new data prediction risk. To calculate it, we independently generate a new dataset $(X_{new}, Y_{new})$ containing $n$ i.i.d. samples drawn according to our data generating mechanism. The prediction risk of the predictor $\widehat{Y}_{new}$ is calculated as $\|\widehat{Y}_{new} - Z_{new}\beta\|^2/n$.

##### 2.3.2 Varying $p$, $K$ and SNR one at a time

To vary $p$ and $K$ one at a time, we first set $n = 300$, $K = 10$, $m = 5$ and choose $p$ from $\{200, 400, 600, 800, 1000\}$, then fix $n = 300$, $p = 600$, $m = 5$ and vary $K$ in $\{10, 20, 30, 40, 50\}$. Both settings use $\zeta = 0.5$. We plot the prediction risks of the four predictors listed above.

To vary the signal-to-noise ratio $\xi = \lambda_K(A\Sigma_Z A^T)/\lambda_1(\Gamma)$, we fix $\Sigma_Z$ and $\Gamma$, and generate $A_J$ by choosing $\zeta \in \{0.1, 0.3, 0.5, 0.7, 0.9, 0.95, 0.99\}$. We set $n = 300$, $p = 400$, $K = 10$ and $m = 3$. For each $\zeta$, we calculate the SNR and plot the prediction risks of each predictor.

**Summary:** Overall, the prediction error for all four methods deteriorates as $K$ increases or the SNR decreases. This indicates that prediction becomes more difficult for large $K$ and small SNR. On the other hand, ER, PFR and Lasso perform better as $p$ increases. This contradicts the classical understanding that having more features increases the degrees of freedom of the model, hence inducing larger variance. By contrast, in our setting, increasing the number of features provides information that can be used to predict $Z$ more accurately. This phenomenon has been observed in the classical factor (regression) model, see, for instance, (5–9).

Among the four candidates, ER has the smallest prediction error in all settings and PLS has the worst performance in most of the settings. Furthermore, PFR fails to detect $K$ and tends to select $\widehat{K} < K$ in the second and third scenarios. It is clear that using $\widehat{K} < K$ leads to a large loss in prediction accuracy. This also indicates that, for principal component regression approaches, detecting $K$ requires larger SNR than making consistent prediction with true $K$ given. In the first plot, we are in a moderate SNR regime and PFR has comparable performance to ER. In the second plot, as $K$ increases, the advantage of ER becomes considerable, which supports the fact that PFR only has guarantees for fixed $K$. Finally, in the third plot, the performance of PFR is more sensitive to the SNR comparing to the other three methods.

### 3 Methodology under CausER

CausER is a novel method that combines the discovery of significant latent factors from Essential Regression with the causal inference implemented in CausalMGM, a method for learning causal graphs over mixed continuous and categorical data (10). This addresses two of the largest challenges of performing causal inference on biological datasets: high dimensionality and multicollinearity among variables. Highly collinear features are grouped into individual latent factors by the LOVE algorithm B, and the significant latent factor selection performed by ER 2.1.2 reduces the dimensionality of the dataset without discarding any latent factors causally linked to the response variable that we wish to predict or make inferences about.

#### 3.1 Methodology under CausalMGM

##### 3.1.1 Mixed Graphical Models

A Mixed Graphical Model (MGM) is an undirected graphical model capable of representing the joint distribution over datasets containing both continuous and categorical variables (11). The model is given by:

$$p(x, y; \theta) \propto \exp\left(\sum_{s=1}^{p}\sum_{t=1}^{p} -\frac{1}{2}\beta_{st}x_s x_t + \sum_{s=1}^{p}\alpha_s x_s + \sum_{s=1}^{p}\sum_{j=1}^{q}\rho_{sj}(y_j)x_s + \sum_{j=1}^{q}\sum_{r=1}^{q}\phi_{rj}(y_r, y_j)\right), \tag{3.1}$$

where $\theta$ represents the full set of parameters, $x_s$ is the $s$th of $p$ continuous variables, and $y_j$ is the $j$th of $q$ categorical variables. The parameter $\beta_{st}$ represents the edge potential between continuous variables $x_s$ and $x_t$, $\alpha_s$ represents the node potential of continuous variable $s$, $\rho_{sj}$ represents the edge potential between continuous variable $x_s$ and categorical variable $y_j$, and $\Phi_{rj}$ represents the edge potential between categorical variables $y_j$ and $y_r$. A non-zero edge potential indicates the presence of an edge in the graph, and thus a pairwise conditional dependence relationship between those two variables. This model has the favorable property that the conditional probabilities of each variable can be represented with a Gaussian linear regression and multinomial logistic regression for continuous and categorical variables respectively. While learning this model over high-dimensional datasets directly is intractable due to the computation of the partition function, the above property enables us to learn the graphical model by minimizing the negative log-psuedolikelihood, given by 3.2, where $\Theta$ refers to the full set of model parameters.

$$\tilde{\ell}(\Theta \mid x, y) = -\sum_{s=1}^{p}\log p(x_s \mid x_{\backslash s}, y; \Theta) - \sum_{j=1}^{q}\log p(y_j \mid x, y_{\backslash j}; \Theta) \tag{3.2}$$

In order to ensure sparsity in the final model, we use proximal gradient descent to fit a penalized version of the negative log-pseudolikelihood from (12), given in 3.3. We identify the optimal values for regularization parameters $\lambda_{CC}$, $\lambda_{CD}$, and $\lambda_{DD}$ using Stable Edge-specific Penalty Selection (StEPS), a method based on model stability, as defined in (12).

$$\underset{\Theta}{\text{minimize}}\, \ell_\lambda(\Theta) = \tilde{\ell}(\Theta) + \lambda_{CC}\sum_{s=1}^{p}\sum_{t=1}^{s-1}|\beta_{st}| + \lambda_{CD}\sum_{s=1}^{p}\sum_{j=1}^{q}||\rho_{sj}||_2 + \lambda_{DD}\sum_{j=1}^{q}\sum_{r=1}^{j-1}||\phi rj||_F \tag{3.3}$$

The undirected MGM does not represent a causal graphical model. However, it does identify pairwise conditional dependence relationships, and the resulting adjacencies are a *superset* of the adjacencies in the underlying causal graph. In the asymptotic sample limit, MGM learns the 'moralized graph', which consists of all edges in the causal DAG as well as additional edges between all spouses (nodes that share the same children) in the causal DAG. This means that the adjacencies learned from MGM can be used as the initial set of adjacencies for constraint-based causal inference methods, rather than a fully connected graph. This reduces the search space of the constraint-based causal inference algorithms, resulting in faster causal inference and fewer errors.

##### 3.1.2 Fast Causal Inference

The FCI algorithm (13) learns a causal partial ancestral graph (PAG) from data that may include latent variables. Similar to the PC algorithm, the FCI algorithm performs conditional independence tests to find the skeleton of the final causal graph and orient the colliders (14). To accommodate the possibility of latent confounders in the causally insufficient case, the FCI algorithm must perform additional conditional independence tests that condition on some non-adjacent variables. This is because in the causally sufficient case with no latent confounders, if two variables are conditionally independent they are independent given some subset of their neighbors in the causal graph. However, this is no longer true in the causally insufficient case. The sets of variables that may cause two adjacent variables to be conditionally independent in the presence of latent confounders is characterized as the Possible D-Sep set. During the Possible D-Sep phase of the FCI algorithm, edges are pruned from the graph if the two variables are conditionally independent given one of the sets of variables in the Possible D-Sep set. Finally, all edges are reoriented to have circle endpoints, and the FCI orientation rules are applied as given by (15). In this paper, we use a version of the FCI orientation rules known as FCI-Max, where the initial collider orientation stage is done based on the separating set with the largest $p$ value, as in (10). This heuristic, based on the observation the $p$ values increase monotonically as the conditional dependence decreases (16), has been shown to considerably improve orientation accuracy over the initial implementation of FCI (10).

Constraint-based causal inference algorithms such as FCI require reliable conditional independence tests, which determine if some variables $X$ and $Y$ are independent given a conditioning set $S$, to learn the causal graphical model. Under the null hypothesis that $X$ and $Y$ are independent given $S$, we expect to see that $P(X \mid Y, S)$ is equal to the null model, $P(X \mid S)$. To accommodate mixed datasets as in (17), we compute these conditional probabilities with either linear regression, in the case that $X$ and $Y$ are continuous, or multinomial logistic regression, in the case that $X$, $Y$, or both are categorical. In the case that the test is performed by a linear regression, we perform a $t$ test on the coefficient of $Y$. If the $p$ value of the test is less than the significance threshold $\alpha$, then we reject the null hypothesis of conditional independence between $X$ and $Y$ given $S$. Alternatively, in the case that the test is performed by a multinomial logistic regression, we perform a likelihood ratio test (LRT) to determine whether $P(X \mid Y, S)$ is equal to the null model, $P(X \mid S)$. Again, if the $p$ value of the test is less than the significance threshold $\alpha$, then we reject the null hypothesis of conditional independence between $X$ and $Y$ given $S$. Note that the conditioning set $S$ can contain both continuous and categorical variables, where categorical variables are transformed into an array of binary indicator variables.

The graphical causal models learned by FCI are PAGs, which are a representation of ancestry in causal graphs that is valid in the presence of latent confounders. Edges in this type of graph have three different types of endpoints: (o, $>$, -), and each represents an ancestral relationship between nodes in the graph. For example, $X \to Y$ indicates that $X$ is an ancestor of $Y$, while $X \leftrightarrow Y$ indicates that a latent confounder causes both $X$ and $Y$. The circular endpoint indicates uncertainty about the true causal endpoint. Thus, $X$ o$\to$ $Y$ could be either $X \to Y$ or $X \leftrightarrow Y$

in the true graph, meaning that the only certain knowledge according to the PAG is that $Y$ is not an ancestor of $X$. An extensive theoretical grounding of these concepts can be found in (14).

###### 3.1.3 Markov blanket

To build a predictor of a given response variable that is informed by the causal structure of variables in the dataset, a small subset of variables known as the Markov blanket are used. This set of variables are the variables that, when conditioned on, make the response variable independent of every other variable in the dataset according to the structure of the causal graph. In the case of a DAG, this set simply contains the parents, children, and spouses (other parents of the children) of the response variable (18). In the case of a PAG, the presence of latent confounders complicates the issue. In addition to the parents, children, and spouses of the response variable, we must include variables linked to the response variable or its children by a latent confounder ($X \leftrightarrow Y$) or a possible latent confounder (*e.g.*, $X \circ\!\!\rightarrow Y$), as well as the parents of those variables. If there is a chain of edges denoting the presence of latent confounders or possible latent confounders (*e.g.*, $W \leftrightarrow X \leftrightarrow Y \leftrightarrow Z$) that is linked to the response variable or its children, then the parents of each node in the chain is included in the Markov blanket (19). However, in practice this can lead to large Markov blankets and a drop in predictive performance, so we set the limit on the number of consecutive variables linked by latent confounders to include to 2. In the previous chain for example, if $W$ was the response variable or one of its children, then $X$, $Y$, and their parents would be included in the Markov blanket.

By this definition, the Markov blanket of a response variable is the minimum set of features that contains all of the information available in the dataset for predicting the response variable. This can result in highly interpretable predictive models with few features and predictive performance that typically matches or exceeds those of models built with the full dataset or variables selected by Lasso. This has previously been successfully applied in biomedical applications, such as predicting whether lung nodules detected in low-dose CT scans are cancerous (20).

##### 3.2 Constructing a causal model on significant latent factors

To identify the latent factors to use in the construction of a causal model, we perform Essential Regression on the variable of interest. In the case that $\widehat{K}$ is small compared to $n$, we learn a causal model over all latent factors identified by LOVE, the response variable, and any categorical variables that we wish to include in the model using CausalMGM. In the case that $\widehat{K}$ is large compared to $n$, we use the Dantzig estimator $\widehat{\beta}_d$ in 2.5 to identify latent factors that are significantly associated with the response variable. We then construct a causal model with the latent factors that have non-zero coefficients in $\widehat{\beta}_d$, the response variable, and any categorical variables we wish to include in the model using CausalMGM.

When constructing the causal model, we first learned an undirected graphical model with MGM 3.1.1 (or GLASSO (21) if fully continuous). The optimal regularization parameters were selected based on graph stability using StEPS (12) (or StARS (22) if fully continuous). The resulting undirected graph was then used as an initial graph for performing causal inference with the FCI algorithm 3.1.2. This yields a causal PAG over the significant latent factors, the response variable, and any additional categorical variables, which can be used for the construction of predictive models or inference about causal mechanisms.

##### 3.3 Stability-based $\alpha$ threshold selection

When building predictive models, the main structural feature of interest in the causal graph is the Markov blanket of the response variable. To select an optimal $\alpha$ threshold value for the conditional independence tests performed by the FCI algorithm, we took a stability-based approach based on StARS. While StARS was originally used for selecting the regularization parameter to be used in GLASSO, we use the same method of subsampling, learning the graph structure, and calculating the instability across subsamples to select an optimal $\alpha$ threshold for learning the Markov blanket of the response variable. However, instead of calculating the instability of an edge in the graph, we calculate the instability of a variable's inclusion in the Markov blanket.

We define $\widehat{\theta}_j(\alpha)$ as the frequency of a variable $j$'s membership in the Markov blanket. Using this, we can calculate instability of a single variable $j$'s membership in the Markov blanket as

$$\widehat{\xi}_j(\alpha) = 2\widehat{\theta}_j(\alpha)\left(1 - \widehat{\theta}_j(\alpha)\right). \tag{3.4}$$

This definition of the instability is twice the variance of the Bernoulli indicator for variable $j$'s membership in the Markov blanket. Additionally, it can be interpreted as the probability of the Markov blankets learned on any two subsamples disagreeing about variable $j$'s membership in the Markov blanket. This arises from the probabilities of the two possibilities for disagreement between subsamples: the probability that the first subsample includes variable $j$ in the Markov blanket and the second subsample excludes variable $j$ can be given as $\widehat{\theta}_j(\alpha)\left(1 - \widehat{\theta}_j(\alpha)\right)$, while the reverse can be given as $\left(1 - \widehat{\theta}_j(\alpha)\right)\widehat{\theta}_j(\alpha)$. When summed, this gives our definition of instability for a single variable $j$'s membership in the Markov blanket. With this, we define the instability of the Markov blanket as a whole, $\widehat{D}(\alpha)$, as

$$\widehat{D}(\alpha) = \frac{\sum_{j \in MB} \widehat{\xi}_j(\alpha)}{m}, \tag{3.5}$$

where $MB$ is the set of variables that shows up in the Markov blanket of the response variable in at least one subsample and at least one value of $\alpha$, and $m$ is the size of set $MB$. Very high values of $\alpha$ will lead to very dense but also very stable graphs, which is undesirable. To avoid this, we monotonize the instability of the Markov blanket as done in StARS, giving

$$\overline{D}(\alpha) = \sup_{0 \leq t \leq \alpha} \widehat{D}(t). \tag{3.6}$$

As it it more difficult to test the instability of many $\alpha$ parameters with CausalMGM and CausER than with the regularization $\lambda$ in GLASSO, we modify the selection of the optimal $\alpha$ threshold $\widehat{\alpha}$. While StARS selects the smallest value of $\lambda$ with an instability less than some threshold $\gamma$, we select the value of $\alpha$ with an instability closest to some threshold $\gamma$, given by

$$\widehat{\alpha} = \inf_{\alpha} |\overline{D}(\alpha) - \gamma|. \tag{3.7}$$

This method for selecting $\widehat{\alpha}$ requires the selection of a threshold $\gamma$. This may make the method seem redundant, as we require a new hyperparameter in order to select $\widehat{\alpha}$. However, this threshold $\gamma$ has a clear interpretation in the context of the learned graph; it represents the average probability of graphs learned on two subsamples disagreeing on the membership of a variable in the Markov blanket. In contrast, the stability of the graph can vary considerably at the same values of $\alpha$ in different datasets. Thus, by setting the threshold $\gamma = 0.05$, we are selecting the value of $\alpha$ that results in Markov blanket selections where the average probability that a variable is present in one selection but not in another is closest to 0.05.

##### 3.4 Causally informed prediction of the response variable

Once the optimal threshold value $\widehat{\alpha}$ is selected with the above method, we build a final causal model using the full dataset and the conditional independence test threshold $\widehat{\alpha}$. We then identify the Markov blanket of the final causal model to be used as predictors in the construction of regression models for the response variable. If the response variable is continuous, we use linear regression, and if the response variable is categorical we use multinomial logistic regression. Predictive performance is estimated using leave-one-out cross-validation, where we train the model on all but one sample and then predict the value of the held out sample. This procedure is repeated for each sample in the dataset, and the performance metric is calculated using the predictions of the held out values.

#### 4 Specifications of the data analysis

We provide detailed specifications of all the data analysis carried out in this paper.

##### 4.1 Imputation of missing values

Among the datasets that we studied, there are different levels of missingness. To impute the missing values, we use the averaged value of the 5 nearest neighbors in Euclidean distance.

##### 4.2 Implementation of different methods

Throughout our analysis, we consider the following competitive methods:

1. CausER: CausER in Section 3.

2. ER: Essential Regression in (2.6) of Section 2.1.

3. CR: Composite Regression in (2.7) of Section 2.1. The tuning parameter of the Lasso step uses the $k$-fold cross validation. We use $k = 10$ if there are more than 3 observations per fold, otherwise set $k = 5$.

4. Lasso: The Lasso (23) from `glmnet` implemented in `R` with the tuning parameter $\lambda_{lasso}$ selected from the $k$-fold cross validation with $k$ chosen according to the same rule for CR. When Lasso selects no feature, we randomly select 5 features and use an ordinary least squares estimator based on these 5 features.

5. PFR: principal factor regression which regresses $\mathbf{Y}$ on the first $K$ principal components of $\mathbf{X}$ where $K$ is selected based on the criterion proposed in (24, 25). Specifically, we estimate $K$ by

$$\widehat{K} = \arg \max_{k \in \{1,2,\ldots,\bar{K}\}} \frac{\widehat{\lambda}_k}{\widehat{\lambda}_{k+1}} \tag{4.1}$$

   where $\widehat{\lambda}_1, \widehat{\lambda}_2, \cdots$ are the non-decreasing eigenvalues of $\mathbf{X}^\top \mathbf{X}/n$ and $\bar{K}$ is some prespecified value, for instance, the largest integer that is no greater than $\min(n, p) - 1$.

6. PLS: partial least squares regression from `plsr` implemented in `R` with the number of components selected by the default function `selectNcomp`.

##### 4.3 Cross-validated assessment of predictive performance

Two cross-validation techniques were used to assess the predictive performance of the different methods: (1) replicated 10-fold cross-validation, and (2) leave-one-out cross-validation.

1. **Replicated 10-fold cross-validation:** For assessing the accuracy of the classifiers in the RTS,S vaccine-induced transcriptomic profiles dataset and the Term / pre-term infants stereotypic immune convergence dataset, 50 replicates of nested 10-fold cross-validation was performed. On each fold, in each replicate, we independently ran each of the methods in 4.2 and assessed the predictive accuracy. For ER, the latent factors were learned on each fold and each replicate, and the regression and final latent factor selection were repeated. For CausER, a causal model was learned over the latent factors selected as significant by ER for each fold and replicate. The average cross-validation accuracy across the 10 folds was calculated for each of the 50 replicates.

2. **Leave-one-out cross-validation:** For all three datasets, we performed leave-one-out cross-validation to assess the accuracy of each method. In leave-one-out cross-validation, each sample in the dataset is held out as the predictive models are trained on the remaining $n-1$ samples, and then the held out sample is predicted with the trained models. The assessment of model performance is done with the set of predictions of the left out values. These predictions were used to calculate Receiver Operating Characteristic (ROC) curves, correlations between predictions and true values, and classification accuracies.

##### 4.4 Multi-omic responses to the Zostavax vaccine dataset

To construct the dataset of multi-omic responses to the Zostavax vaccine, we included the following multi-scale measurements of immune state: IgG titers, blood transcriptional modules, metabolic clusters, $CD4^+$ T cell populations, $T_{FH}$ cell populations, flow cytometry cell populations, cytokine profiles, and IFN$\gamma$ T cells. We used subject age as the response variable for the n = 72 subjects. We exclude the features that have missing values for more than a half of subjects. We also exclude 5 subjects that have no observed features. The remaining data sets are merged via the unique id's of subjects. The final data set contains $p = 1721$ features of $n = 67$ subjects.

We applied ER with $\delta = 0.38$ and obtained $\widehat{K} = 57$ clusters. As $\widehat{K}$ is relative large comparing to $n$, we used the Dantzig estimator $\widehat{\beta}_d$ of $\beta$ in (2.5) and the non-zero support $\widehat{\beta}_d$ selects 18 significant factors for predicting the response.

For this dataset, which is fully continuous, the initial undirected skeleton was learned with Graphical LASSO (GLASSO) (21) implemented in the `huge` package in `R` (26). The optimal regularization parameter $\lambda = 0.32$ was selected by StARS (22). The causal model over the latent factors was built using FCI-Max, as implemented in the `rCausalMGM` package in `R` (`https://github.com/tyler-lovelace1/rCausalMGM`). The optimal conditional independence test significance threshold for learning the causal graph, $\alpha = 0.1$, was selected as described in 3.3.

##### 4.5 Term / pre-term infants stereotypic immune convergence dataset

For pre-processing the data, we first combine the data sets of Cell population frequencies and Short final ComBat by removing the irrelevant features such as "gender", "mode of delivery", "family" etc. Then we further exclude the control samples with row indices from 326 to 337. We further pulled out the subdata with "Relation" equal to "child" and the final data set we use has $n = 183$ samples and $p = 282$ features with 56 samples from week 1 and 46 samples from week

12. The response is binary, either "Control" (representing term) or "Premature" (representing pre-term). We use the 5-NN to impute the missing values.

We perform the classification of pre-term / term by using the features collected in week 12. We applied ER with $\delta = 0.15$ and obtained 14 clusters. Our estimator $\widehat{\beta}$ is constructed via (2.3) and the 95% confidence intervals select two significant factors $Z_5$ and $Z_7$ for predicting term and pre-term.

For features in week 1, we used $\delta = 0.17$ with 14 clusters and the significant factors include $Z_3$, $Z_4$, $Z_{10}$, $Z_{11}$ and $Z_{14}$.

Only week 12 data was analyzed with CausER. For this dataset, only two significant latent factors, $Z_5$ and $Z_7$, were identified, making causal orientations unidentifiable in most cases (the only exception being the graph $Z_5 \circ\!\!\rightarrow Y \leftarrow\!\!\circ Z_7$). Additionally, there are too few features for stability-based selection of $\alpha$. However, a causal model was constructed over all latent factors using FCI-Max, as implemented in the `rCausalMGM` package in `R` (https://github.com/tyler-lovelace1/rCausalMGM). The optimal conditional independence test significance threshold for learning the causal graph, $\alpha = 0.2$, was selected as described in 3.3.

##### 4.6 RTS,S vaccine-induced transcriptomic profiles dataset

By concatenating the two gene-expression data sets, we end up with $n = 116$ samples with $p = 22277$ probes. We filtered out the probes that could map to multiple genes, and then the technical replicates were averaged with the `limma` package in `R` (27), giving the expression of $p = 12424$ genes.

The responses $Y \in \mathbb{R}^n$ are categorical representing three time points. We applied ER to the data set with selected $\delta = 0.04$ and obtained $\widehat{K} = 1674$ clusters. The estimator of $\beta$ is the Dantzig estimator in (2.5) which has 86 non-zero elements. This implies there are 86 significant factors $Z$ for predicting the response.

For this dataset, which is mixed, the initial undirected skeleton was learned with MGM, described in 3.1.1, implemented in the `rCausalMGM` package in `R`. The optimal regularization parameters $\lambda_{CC} = 0.27$, $\lambda_{CD} = 0.27$ ($\lambda_{DD}$ was irrelevant because there is only one categorical variable) were selected by StEPS (12). The causal model over the latent factors was built using FCI-Max, as implemented in the `rCausalMGM` package in `R` (https://github.com/tyler-lovelace1/rCausalMGM). The optimal conditional independence test significance threshold for learning the causal graph, $\alpha = 0.2$, was selected as described in 3.3.

### A Identifiability results of $A$ and $\beta$

We re-state the identifiability results of $A$ and $\beta$ from (1) and (2).

**Theorem 1** (Theorems 1 & 2 (1)). *Under model $X = AZ + E$ with Assumption 1, the set of pure variable $I$, its partition $\mathcal{I} = \{I_1, \ldots, I_K\}$*[2] *and the number of factors are identifiable from $\Sigma = Cov(X)$.*

*Moreover, the matrix $A$ is identifiable up to a $K \times K$ signed permutation matrix.*

**Proposition 2** (Proposition 1 (2)). *Under model (1.1) – (1.2) with Assumption 1, the coefficient vector $\beta$ is identifiable up to a signed permutation matrix.*

### B The LOVE algorithm

We first give the specifics of estimating $I$ and $K$ developed by (1).

---

**Algorithm 1** Estimate the partition of the pure variables $\mathcal{I}$ by $\widehat{\mathcal{I}}$

---

**procedure** PureVar($\widehat{\Sigma}$, $\delta$)
    $\widehat{\mathcal{I}} \leftarrow \varnothing$.
    **for all** $i \in [p]$ **do**
        $\widehat{I}^{(i)} \leftarrow \left\{ l \in [p] \setminus \{i\} : \max_{j \in [p] \setminus \{i\}} |\widehat{\Sigma}_{ij}| \le |\widehat{\Sigma}_{il}| + 2\delta \right\}$
        $Pure(i) \leftarrow True.$
        **for all** $j \in \widehat{I}^{(i)}$ **do**
            **if** $\left| |\widehat{\Sigma}_{ij}| - \max_{k \in [p] \setminus \{j\}} |\widehat{\Sigma}_{jk}| \right| > 2\delta$ **then**
                $Pure(i) \leftarrow False,$
                **break**
        **if** $Pure(i)$ **then**
            $\widehat{I}^{(i)} \leftarrow \widehat{I}^{(i)} \cup \{i\}$
            $\widehat{\mathcal{I}} \leftarrow$ Merge($\widehat{I}^{(i)}$, $\widehat{\mathcal{I}}$)
    **return** $\widehat{\mathcal{I}}$ and $\widehat{K}$ as the number of sets in $\widehat{\mathcal{I}}$

**function** Merge($\widehat{I}^{(i)}$, $\widehat{\mathcal{I}}$)
    **for all** $G \in \widehat{\mathcal{I}}$ **do** ▷ $\widehat{\mathcal{I}}$ is a collection of sets
        **if** $G \cap \widehat{I}^{(i)} \ne \varnothing$ **then**
            $G \leftarrow G \cap \widehat{I}^{(i)}$ ▷ Replace $G \in \widehat{\mathcal{I}}$ by $G \cap \widehat{I}^{(i)}$
            **return** $\widehat{\mathcal{I}}$
    $\widehat{I}^{(i)} \in \widehat{\mathcal{I}}$ ▷ add $\widehat{I}^{(i)}$ in $\widehat{\mathcal{I}}$
    **return** $\widehat{\mathcal{I}}$

---

Next, for each $a \in [\widehat{K}]$ and $b \in [\widehat{K}] \setminus \{a\}$, we compute

$$\left[\widehat{\Sigma}_Z\right]_{aa} = \frac{1}{|\widehat{I}_a|(|\widehat{I}_a| - 1)} \sum_{i,j \in \widehat{I}_a, i \ne j} |\widehat{\Sigma}_{ij}|, \qquad \left[\widehat{\Sigma}_Z\right]_{ab} = \frac{1}{|\widehat{I}_a||\widehat{I}_b|} \sum_{i \in \widehat{I}_a, j \in \widehat{I}_b} \widehat{A}_{ia}\widehat{A}_{ib}\widehat{\Sigma}_{ij}, \tag{B.1}$$

to form the estimator $\widehat{\Sigma}_Z$ of $\Sigma_Z$. Furthermore, we restate the estimation of $A_{I\cdot}$ in (1). For each $k \in [\widehat{K}]$ and the estimated pure variable set $\widehat{I}_k$,

$$\text{Pick an element } i \in \widehat{I}_k \text{ at random, and set } \widehat{A}_{i\cdot} = e_k; \tag{B.2}$$

$$\text{For the remaining } j \in \widehat{I}_k \setminus \{i\}, \text{ set } \widehat{A}_{j\cdot} = \text{sign}(\widehat{\Sigma}_{ij}) \cdot e_k. \tag{B.3}$$

[2] $\mathcal{I}$ is identifiable up to a group permutation.

For the estimation of $A_{J\cdot}$, we use the Dantzig-type estimator $\widehat{A}_D$ proposed in (1) given by

$$\widehat{A}_{j\cdot} = \arg\min_{\beta^j} \left\{ \|\beta^j\|_1 : \left\| \widehat{\Sigma}_Z \beta^j - (\widehat{A}_{\widehat{I}\cdot}^\top \widehat{A}_{\widehat{I}\cdot})^{-1} \widehat{A}_{\widehat{I}\cdot}^\top \widehat{\Sigma}_{\widehat{I}j} \right\|_\infty \leq c\sqrt{\log(p \vee n)/n} \right\} \tag{B.4}$$

for any $j \in \widehat{J}$, with some constant $c > 0$. The estimator $\widehat{A}$ enjoys the optimal convergence rate of $\max_{j\in[p]} \|\widehat{A}_{j\cdot} - A_{j\cdot}\|_q$ for any $1 \leq q \leq \infty$ (1, Theorem 5).
