## Supplementary Note 2 for "Essential Regression - a generalizable framework for inferring causal latent factors from multi-omic human datasets"

### Supplementary Note 2 – Applying ER to datasets generated using older technologies

#### Analyzing latent factors potentially reflective of trained immunity in a vaccine response

To test whether ER is applicable to datasets generated using older technologies, we applied to analyze the temporal dynamics of transcriptional responses (microarray data) induced by the malaria RTS,S vaccine(1). RTS,S has a standard regimen of 3 doses separated by a month, and is currently the most advanced malaria vaccine candidate, that has consistently demonstrated 40-80% protective efficacy in malaria-naïve individuals in controlled human challenge studies(2). There has been intense interest over the last decade at uncovering molecular signatures induced by the RTS,S vaccine and corresponding correlates of protection (2-4). In a controlled human infection setting, differential expression of immunoproteasome genes was identified as a pre-challenge correlate of protection (1). After the third dose, as expected, there was a striking but transitory shift in inflammatory gene expression followed a convergence of the majority of gene signatures back to pre-vaccination levels within 2 weeks after the third dose (1). We reasoned that aspects of trained immunity induced by the vaccine may be reflected in the transcriptomic signatures that do not converge after 2 weeks. Thus, a sensitive method such as ER would be able to discriminate between expression profiles at the following time-points – pre-vaccination (G1), the day after the third dose (G2) and 14 days after the third dose (G3) (Fig. S3a) and reveal candidate genes and molecular pathways that could contribute to trained immunity. In this instance, the use of a microarray dataset also afforded the opportunity to explore how ER performs with noisier but nevertheless valuable datasets generated using older technologies.

As before, the ability of the different methods to discriminate between G1, G2 and G3 transcriptional profiles was measured in a rigorous cross-validation framework (Methods). We found that there were significant differences in the ability of the different methods to discriminate between the three kinds of expression profiles, with ER and CausER (CausalMGM on the significant latent factors from ER) having the best performance, significantly better than the other methods ($P < 0.01$, Figs. S3b, S3c). Next, we chose to focus on the ability of the different methods to specifically distinguish the G3 profile from the other two (Fig. S3d) or just the G1 profile (Fig. S3e). This constituted the most “difficult” discrimination as there are broad differences in the expression profiles between the pre- (G1) and 24-hour-post-vaccination (G2) time-points, but most of these differences disappear by 14 days (G3) (1). Consistent with expectation, in this binary classification setting, there was wide variability in the performance of the methods to specifically discriminate the G3 time-point from the G1 and G2 time-points. While PFR and PLS performed poorly, CausER, ER and LASSO had significantly better performance, with CausER being the best performing method ($P < 0.01$, Figs. S3d and S3e). In terms of correctly classifying just the true G3 profiles as G3, PLS and PFR had poor performance, while CausER had the best performance, significantly better than other methods ($P< 0.01$, Fig. S3f).

Next, we focused on the CausER hits i.e., the significant latent factors from ER in the Markov blanket of the outcome variable (Fig. S3g). Genes comprising these latent factors were seen to be differentially expressed between the G1 and G3 samples (Fig. S3h, Fig. S3i). Our results suggest that beyond the initial divergence of immunoproteasome genes, there is a sustained divergence (2 weeks post-vaccination) of genes involved in immune-metabolic processes. These results complement recent findings that suggest that targeting immunometabolism is a promising direction in modulating trained immunity (5). While a vaccine induces a rapid initial divergence in inflammatory signatures reflecting the activation of innate immune cells and their engagement with adaptive B and T cells, it may also induce alterations in the innate immune compartment that are discernible at later time points and contribute to a distinct form of immune memory (5).
